# Glycosylation and Crowded Membrane Effects on Influenza Neuraminidase Stability and Dynamics

**DOI:** 10.1101/2023.09.10.556910

**Authors:** Christian Seitz, İlker Deveci, J. Andrew McCammon

**Author notes:** ^1^Department of Computer Science, University of Chicago, Chicago, Illinois and ^2^Data Science and Learning Division, Argonne National Laboratory, Chicago, Illinois (Christian Seitz).

## Abstract

All protein simulations are conducted with varying degrees of simplifications, oftentimes with unknown ramifications on how these simplifications affect the interpretability of the results. In this work we investigated how protein glycosylation and lateral crowding effects modulate an array of properties characterizing the stability and dynamics of influenza neuraminidase. We constructed three systems: 1) Glycosylated neuraminidase in a whole virion (i.e. crowded membrane) environment 2) Glycosylated neuraminidase in its own lipid bilayer 3) Unglycosylated neuraminidase in its own lipid bilayer. We saw that glycans tend to stabilize the protein structure and reduce its conformational flexibility while restricting solvent movement. Conversely, a crowded membrane environment encouraged exploration of the free energy landscape and a large scale conformational change while making the protein structure more compact. Understanding these effects informs what factors one must consider while attempting to recapture the desired level of physical accuracy.

**TOC figure:** TOC. Schematic of the three systems used here. (A) 2009-H1N1-ungly is the unglycosylated, single protein NA system. (B) 2009-H1N1-gly is the glycosylated, single protein NA system. (C) 2009-H1N1-vir is the NA in the virion context.

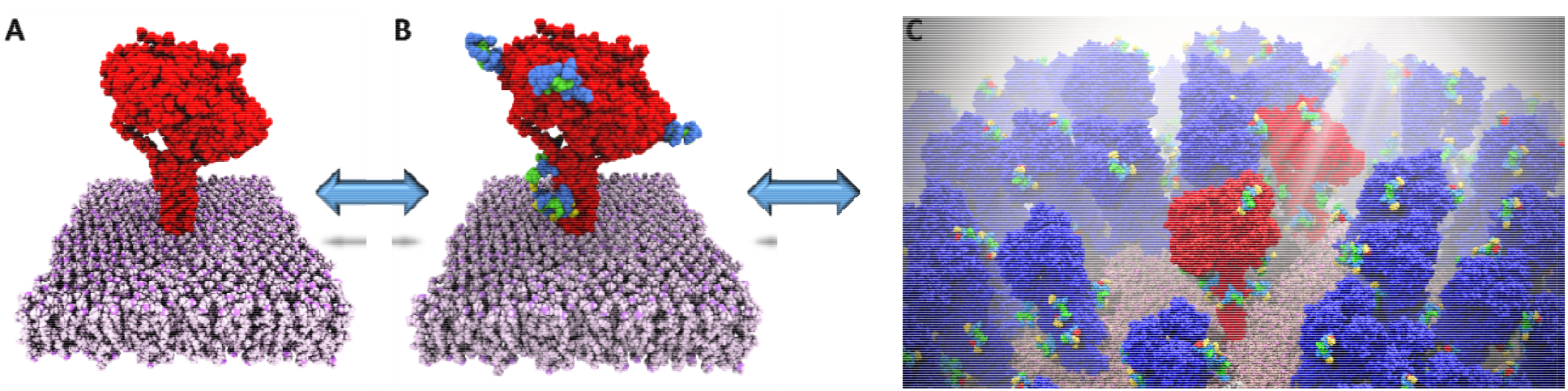

---

Protein molecular dynamics (MD) simulations has been an exciting field ever since the seminal start in 1977. (1) However, many of these simulations have been run on single proteins in solvent without any other complicating factors in play to simplify system setup and analysis. (2) Out of these factors, we selected glycosylation and the protein environment for further study, with the larger goal of informing when one can simplify these aspects to see a given observable and when one cannot. Glycans are present on about half of all proteins (3) and play a diverse role biologically, being implicated in molecular recognition, (4) antibody/drug shielding, (5-7) protein folding, (8,9) intracellular transport, (10) protein clearance, (11) and more. Glycosylation has been shown to increase protein stability using a variety of experimental and computational techniques (9,12-20) but these findings have not been replicated in all protein systems. (21,22) Decoding the effect glycans have on protein dynamics is similarly ambiguous. Some work has shown that glycosylation may decrease the root mean square fluctuations (RMSF), (23-25) increase protein dynamics and flexibility, (22) reduce global protein mobility, (15,26) or that the dynamics of some regions can be dampened by glycosylation while the dynamics of other regions are promoted by glycosylation (27) which can be independent of whether glycans are even found in those regions. (24,28,29)

Environment effects on proteins are also poorly resolved. Proteins are capable of crowding together very effectively (30) and this crowding may stabilize protein structures (31-37) but this stabilization effect is dependent on the protein sequence and its shape (38) which is complicated by the fact that the protein shape itself can be modified by crowding. (39-41) However, there is substantial conflicting literature suggesting that protein crowding destabilizes proteins (42-44) or does not affect their stability at all, (45) suggesting that whether the environment stabilizes or destabilizes a protein is due to its interactions with the environment (42,43,46-52) from differences in enthalpic and entropic interactions. (48,53) Even examining a single protein of interest, some crowding agents can stabilize it (54) while others destabilize it. (55) Even less studied is how protein crowding effects the solvent environment. One study showed how water diffusion is faster in systems with more components (56) which was contradicted by a different study showing how crowding can reduce the solvent dielectric constant and water self-diffusion (57) and that more crowded systems slow water diffusion more than less crowded systems. (58) There is ample room left to investigate how these crowding effects influence protein stability and dynamics. (2) We shed light on this aspect in this work, investigating how glycosylation and a crowded membrane affects protein stability, protein dynamics, and solvent behavior.

We examined three different systems and will refer to them by their abbreviated form throughout the manuscript: a single unglycosylated neuraminidase (NA) tetramer alone in a lipid bilayer, with three replicates of this system simulated (2009-H1N1-ungly), a single glycosylated NA tetramer alone in a lipid bilayer, with three replicates of this system simulated (2009-H1N1-gly), and a single glycosylated NA tetramer in a virus shell with numerous neighboring membrane proteins, with one replicate simulated (2009-H1N1-vir); a representative NA structure is shown in Figure 1 while glycosylation sites are displayed in Table SI1. The 2009-H1N1-gly and 2009-H1N1-ungly systems were constructed and simulated for this work, while the 2009-H1N1-vir system was constructed and simulated previously (59) and is being further analyzed here. Throughout this work, we will refer to “lateral crowding” or a “crowded membrane” interchangeably; here we define this to mean the 2009-H1N1-vir environment, focusing on one membrane protein of interest within a model membrane filled with other membrane proteins.

**Figure 1.**
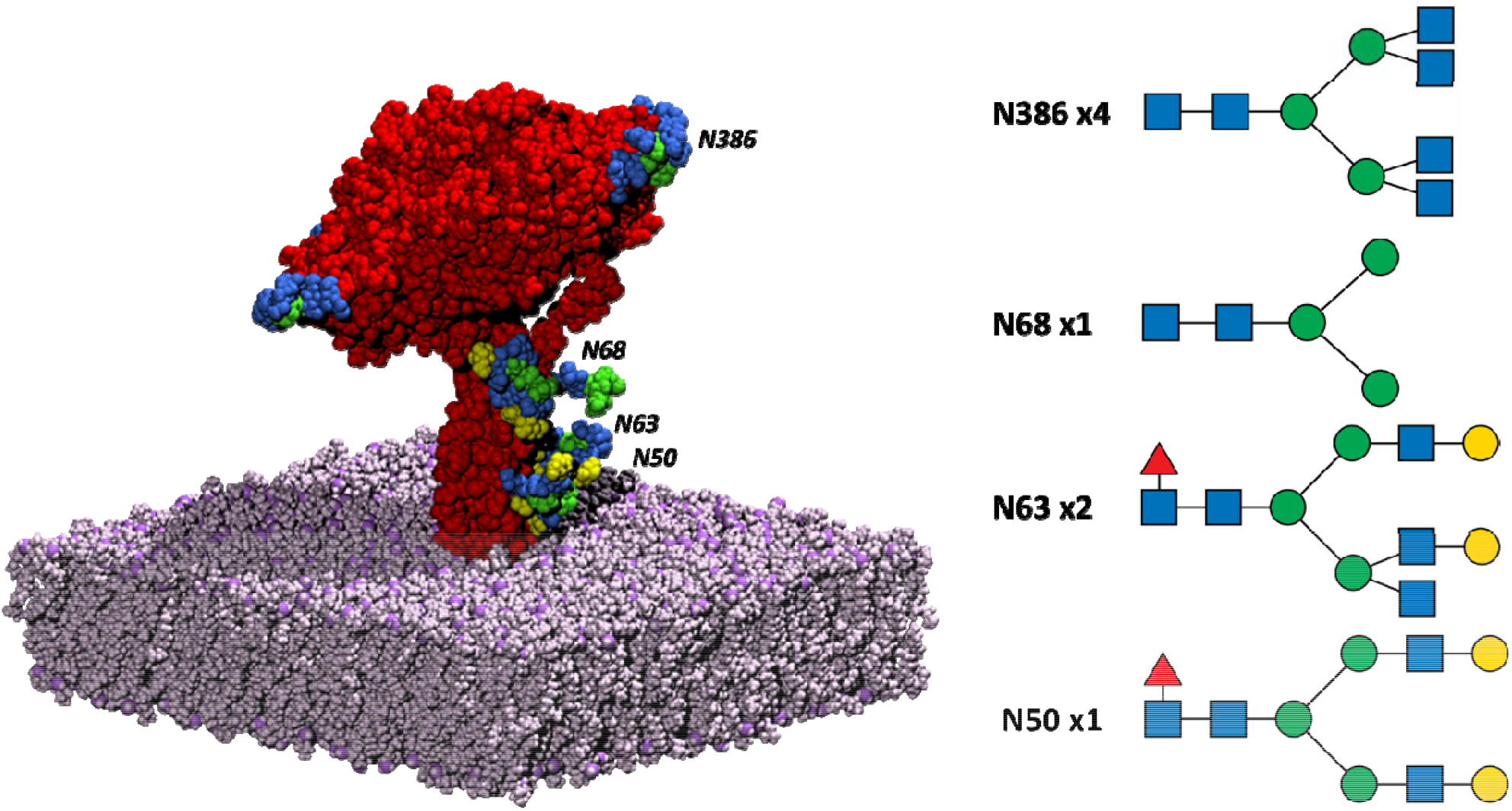
The NA system. The NA tetramer is in red while the bilayer is in mauve. The glycosylation sites and their glycan structures are also shown; the blue squares represent N-acetylglucosamine, the green circles represent mannose, the red triangles represent fucose while the yellow circles represent galactose. Remembering that there are four monomers in the NA tetramer, there is a glycan at the N386 position in each monomer, with two monomers containing a glycan at N68 and one monomer containing a glycan at N68 and N50. The glycan structures were rendered with GlycoGlyph. (60)

Glycans are chains of atoms that attach to the sides of proteins; intuitively this should modify structural stability of the protein, either positively or negatively. This is where our investigations will begin. As a proxy for global protein rigidity, we measure nonpolar interatomic interactions and calculate the energy at which these interactions are excluded; more rigid proteins will retain nonpolar interactions at higher energy cutoffs (see methods in SI (24,61)). We also measure protein compactness through the radius of gyration, and stability through root mean square deviations (RMSDs). The 2009-H1N1-ungly system is slightly more stable at low energy cutoffs — (Figure 2A) due to the flexible glycans being excluded (data not shown). As the energy cutoff is increased —, the 2009-H1N1-gly system becomes a more stable, rigid structure (Figure 2A). These trends are seen whether rigidity is studied as a function of α-carbons or by using every atom in the system (Figure SI1A). Glycosylation increasing protein rigidity has been seen before. (24) When we compare the environment’s effects on rigidity using RMSD clusters of the simulation frames, we see that the 2009-H1N1-vir system is much more rigid when compared to the single protein systems (Figure 2B), while the 2009-H1N1-ungly system is slightly more rigid than the 2009-H1N1-gly system (Figure 2B and Table SI2). We also see a reduction in the radius of gyration *R*_*g*_ of the proteins due to glycosylation, and a further reduction due to a laterally crowded environment (Figure 2C), meaning that they occupy a more compact volume. Previous literature has been divided on this topic: some work shows glycans making proteins more compact (24,25) while other work shows glycans do not affect the compactness of proteins (29) (and personal correspondence (28)). Glycosylation does not affect the NA stalk stability (Figure 2D) but does stabilize the NA head, as measured by RMSD (Figure 2E). Our data cannot discern whether this stabilization occurs due to glycans stabilizing the NA head at every time point, or whether glycans simply increase the time it takes for NA head stabilization to converge. We do not see the same effect for the NA stalk; this may be because the stalk is already less stable so any differences are washed out in the higher RMSD values. In summary, we see that glycosylation increases the rigidity of NA, makes the NA structure more compact, and stabilizes the NA head (Figure 2). Other work has shown that glycans can stabilize the protein structure (23,25,62) (and Figure SI2), do not cause any change in stability, (28,29,63) or destabilize the structure. (64) Similar to what we see, some previous work also shows that glycosylation effects on stability may not be consistent throughout the protein of interest, and may change depending on which sequon is examined. (65)

**Figure 2.**
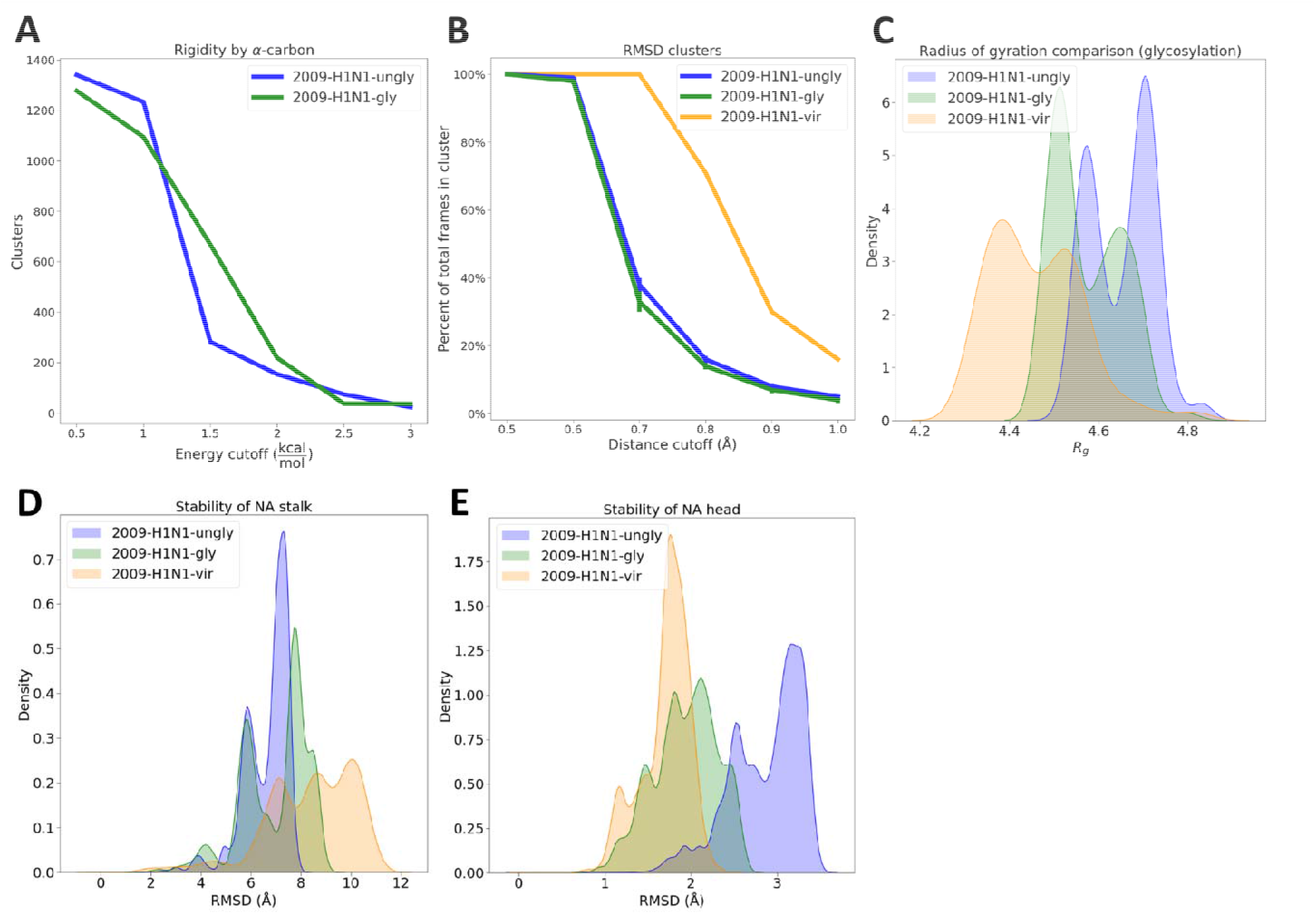
Protein rigidity, compactness and stability as a function of glycosylation and environment. (A) Glycosylation effects on global protein rigidity, showing the number of clusters that retain nonbonded interactions at different energy cutoffs. (B) RMSD clusters of the simulation frames, showing how many clusters are found as a percentage of total frames at each distance cutoff. (C) Radius of gyration of deglycosylated systems. Here, we simulated the 2009-H1N1-gly and 2009-H1N1-vir systems with glycans before removing these glycans for the *R*_*g*_ calculation. Figure SI1B shows the *R*_*g*_ as a function of time. In Figure 3B and Figure SI5B we show the *R*_*g*_ including the glycans. (D) Stability of the NA stalk as a function of the system. Figure SI1C shows the stalk RMSD as a function of time. (E) Stability of the NA head as a function of the system. Figure SI1D shows the head RMSD as a function of time.

Considering that proteins natively exist in a crowded, chaotic environment, much work has gone into investigating how different crowding conditions, and different crowders themselves, influence protein stability. There is a much smaller body of literature examining how a crowded membrane, or “lateral crowding” can affect a membrane protein of interest. (66) In terms of stability, it does not appear that these membrane protein neighbors influence the stability of NA. They slightly destabilize the NA stalk (Figure 2D) but do not affect the NA head (Figure 2E), nor do they affect protein stiffness (Figure 2B). More differences were seen when investigating protein dynamics in a crowded membrane environment.

After our initial tests on rigidity (Figure 2A and Figure 2B), we explored protein rigidity further through internal force constants (eigenvalues) and liquid-solid character (Table 1), where higher eigenvalues correlate to stiffer systems and lower Lindemann values correlate to a more solid system. The internal force constants were measured through three methods: 1) Principal component analysis (PCA) reduces the system into a linear combination of coordinates describing the full system dynamics; 2) Gaussian network models (GNMs) coarse grain the system into α-carbons connected with an interatomic potential to describe isotropic fluctuations (i.e. independent of the direction of measurement); 3) Anisotropic network models (ANMs) also coarse grain the system into α-carbons, this time assuming directional dependence (anisotropy) while the Lindemann coefficient was calculated from the PCA space (see methods in SI (67-75)). Measuring protein stiffness through GNMs and ANMs do not show significant differences due to glycosylation, but an analysis of the principal component space does show that glycans increased the stiffness of NA (Table 1). Considering that stiffer materials will behave more like solids while softer materials will behave more like liquids, we measured the Lindemann coefficient for each of our systems (Table 1). Previous work has shown how phosphorylation can modulate the liquid-solid character of a protein, (76) but we do not see any significant differences due to glycosylation or the environment (Table 1).

**Table 1.** Stiffness and solidity of NA systems. Eigenvalues of the first principal component from PCA and the lowest frequency mode from the GNM and ANM models are shown, along with the Lindemann coefficient. We did not calculate GNM and ANM modes for the 2009-H1N1-vir system because these would be identical to the 2009-H1N1-gly system. The values for the GNM and ANM modes are in arbitrary or relative units, and the Lindemann coefficient does not have a unit.

|  | PC1 | GNM | ANM | Lindemann |
| --- | --- | --- | --- | --- |
| 2009-H1N1-ungly | 4080 Å<sup>2</sup> | 0.0261 | 0.00114 | 0.431 ± 0.060 |
| 2009-H1N1-gly | 5480 Å<sup>2</sup> | 0.0261 | 0.00114 | 0.433 ± 0.049 |
| 2009-H1N1-vir | 4070 Å<sup>2</sup> | N/A | N/A | 0.417 |

After seeing that glycans stabilize the NA head (Figure 2E), we wanted to know if these head glycans would affect NA dynamics. We studied this through fluctuations through the course of the simulations, the solvent accessible surface area (SASA), and two studies on entropy: packing entropy, derived from the volume an amino acid occupies divided by its available volume calculated from a static structure, and dihedral entropy, which uses a one-dimensional approximation of the classical coordinate dihedral entropy calculated from the system trajectories (see methods in SI (77,78)). Due to limitations in sampling, most of these results did not reach the level of statistical significance across our systems. However, considering we observed the same trends across disparate, unrelated techniques, we will discuss the results here. Similar to previous work (23-25,79-81) we saw that glycosylation reduces fluctuations of our protein (Figure SI3A). Interestingly, we show a distance dependence of this effect: The dynamics reduction is stronger the closer one gets to the sequon. Other literature, however, has shown glycosylation can, to a small degree, increase protein dynamics and flexibility (22) or that the dynamics of some regions can be dampened by glycosylation while the dynamics of other regions are promoted by glycosylation. (27) We also examined how glycosylation affects entropic properties of the protein; intuitively one would presume that a reduction in dynamics would be accompanied by a reduction in microstates accessed (and thus a reduction in entropy). This is what we saw when examining packing entropy (Figure SI3B) and dihedral entropy (Figure SI3D and SI3E) – both are reduced due to glycosylation, and both reductions are stronger at the sequon than in the protein as a whole, in agreement with the RMSF work (Figure SI3A). Previous work has shown that proteins with lower entropy will have lower solvent accessibility. (82-84) Despite the 2009-H1N1-ungly system having lower solvent accessibility than the 2009-H1N1-gly system (Figure SI3C), there are no global trends in entropy between these two: the 2009-H1N1-ungly system as a whole has slightly higher φ-angle entropy (Figure SI3D) and ψ-angle entropy (Figure SI3E), while they have roughly the same packing entropy (Figure SI3B).

If glycosylation reduces conformational entropy, does the protein environment affect conformational entropy? We also saw that lateral crowding increases the dihedral entropy of the sequon, but that this did not appear to affect the dihedral entropy throughout the protein as a whole; in fact, it appears that a crowded protein membrane environment slightly decreases dihedral entropy throughout the protein (Figure SI4). In other words, having a crowded membrane environment increases the conformational entropy of the N-linked sequons while decreasing the conformational entropy of the protein as a whole.

When examining large scale simulation work, such as work on viral shell simulations, (59) a persistent question appears: Can we see those same conformational changes and protein characteristics in a simulation with reduced computational cost? We begin to address that significant question here. Some recent work has shown that proteins simulated in a crowded environment can show new motions that are not seen either when the same proteins are simulated individually (59,85) or when they are simulated in an uncrowded environment (86) but transitions between these conformations appear to be slower than in uncrowded environments when they occur. (2) Other work has seen that protein conformational transitions are not correlated with the protein-protein contacts created in a crowded membrane environment. (59) We explore how a crowded membrane affects transitions through NA head tilting, a significant conformational change where the NA head swivels on its stalk (see methods in SI (59)). We see that NA head tilting can be seen in single protein simulations and that this rate of transition is increased by glycans and by using a crowded membrane environment (Figure 3A), although this does not appear to be due to the protein-protein contacts seen in a crowded membrane environment. (59) It may be due to other environmental differences in our systems, or it may simply be a sampling artifact. Regardless, this rate may be slowed down again if a truly crowded environment with free floating proteins were used as the excluded volume effect will come into play. (87) In line with this, some other studies have seen suppressed conformational dynamics due to crowding, (52,88,89) with the caveat that these studies did not examine a membrane crowded with neighboring membrane proteins.

**Figure 3.**
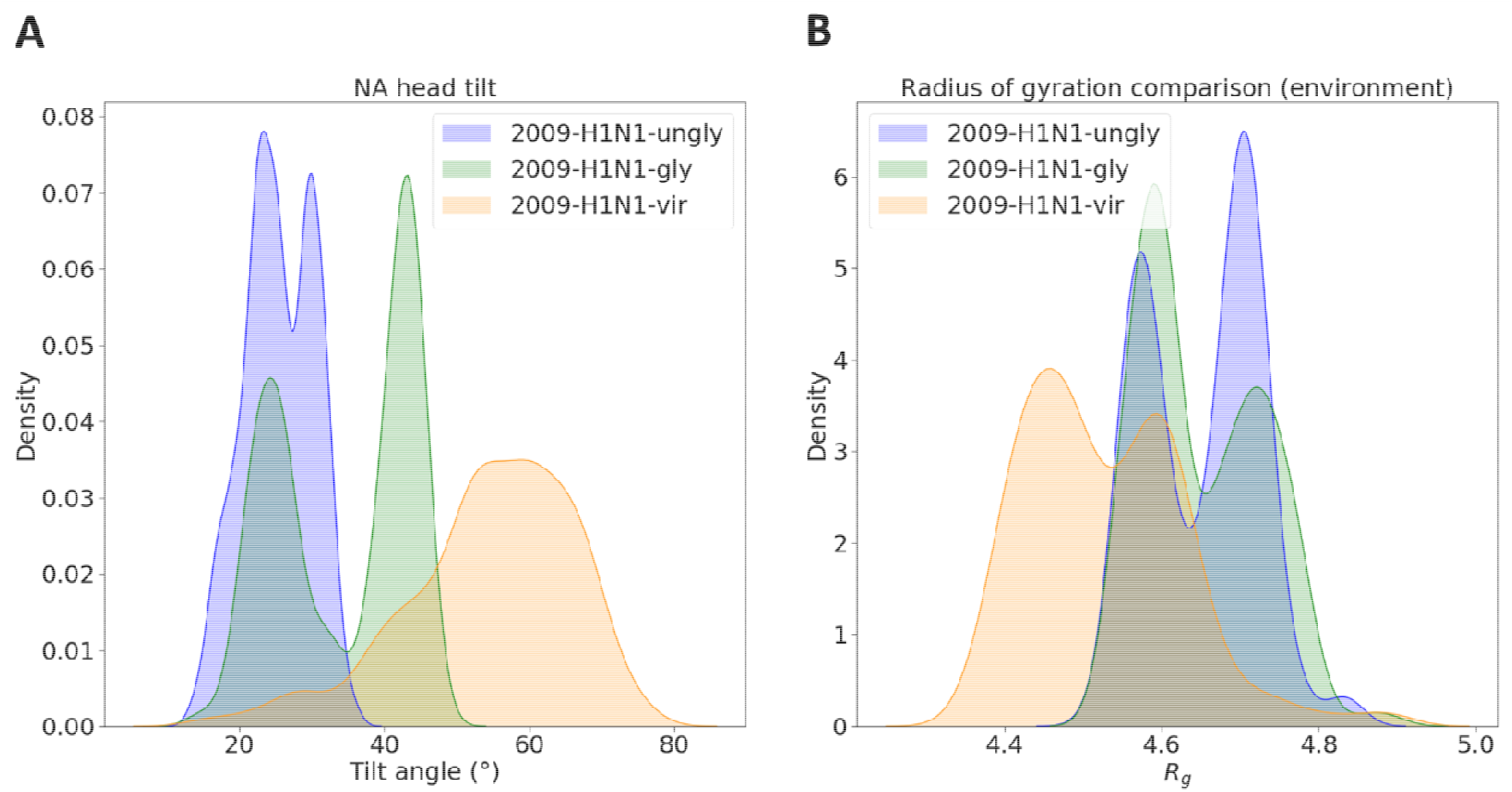
Conformational flexibility and compactness of the NA system. (A) NA head tilt angles. The angle of how much the NA head tilts relative to the stalk was measured. Figure SI5A shows the tilt angle as a function of time. (B) *R*_*g*_ values for NA. Here, we simulated the 2009-H1N1-gly and 2009-H1N1-vir systems with glycans and retained the glycans in the *R*_*g*_ calculation. Figure SI5B shows the *R*_*g*_ as a function of time. In Figure 2C and Figure SI1B we show the *R*_*g*_ after removing the glycans.

The 2009-H1N1-vir system is more compact and has a smaller radius of gyration as a whole than the 2009-H1N1-gly system (Figure 3B) due to the presence of protein neighbors; most literature has seen a crowded environment making a protein more compact (34,39,40,90-92) while two other studies saw no real difference. (45,93) A more compact protein would indicate lower entropy; our 2009-H1N1-vir system as a whole contains less dihedral entropy than the 2009-H1N1-gly system (Figure SI4) whereas a system with higher entropy will have higher solvent accessibility. (82-84) We also saw that our 2009-H1N1-gly system has more dihedral entropy across the protein as a whole than the 2009-H1N1-vir system (Figure SI4) along with higher solvent accessibility (Figure SI3C). In addition, a more compact protein would indicate faster diffusion. This is because *R*_*g*_ is proportional to a protein’s instantaneous diffusivity: a higher *R*_*g*_ means slower diffusion, and vice versa. (94)

Finally, we explored how glycosylation affects sampled space and the harmonic well shape of the first principal component (PC1) from PCA (see methods in SI (79,95-97)). One previous study found that a protein accesses a smaller amount of conformational space in a crowded environment. (79) Conversely, we find that lateral crowding increases the amount of first principal component space NA accessed, with glycans reducing this space accessed (Table 2). This is in line with our RMSF data showing that glycans reduce global protein range of motion (Figure SI3A). Interestingly, the increase in the PC1 space sampled in the 2009-H1N1-vir system as compared to the 2009-H1N1-gly system does not correspond to significant differences in the steepness of the harmonic wells corresponding to PC1 (Table 2). Glycans make NA’s PC1’s harmonic well more shallow and smooth (Table 2) which may help explain why the 2009-H1N1-gly system displays a larger head tilt than the 2009-H1N1-ungly system. We see glycosylation reducing the PC1 space that NA explores (Table 2); glycosylation reducing conformational space sampled was also seen in a study utilizing 500 ns of MD sampling (28), comparable to our 441 ns of sampling, although a different study utilizing 75+ μs of MD sampling saw that glycosylation does not affect the conformational space that one protein of interest can access. (98) This suggests that glycosylation may slow the exploration of the PC1/free energy, but that an exhaustive sampling regime may find similar spaces explored irrespective of glycosylation.

**Table 2.** Internal dynamics of the NA systems as measured through PCA. The 2009-H1N1-ungly system was set as the reference space sampled and the sampling of the other two systems were measured relative to this. *k* is the harmonic force constant; larger *k* values indicate deeper and sharper energy wells explored by the systems, while the internal friction coefficient ζ indicates diffusion through the free energy surface. Larger ζ values indicate more friction, i.e. a rougher potential energy surface, which equates to slower diffusion through that surface. The *α* and *τ* parameters are included for completeness as they are needed for full computation.

| | PC1 space sampled | $k \left( \frac{kJ}{mol \cdot nm^2} \right)$ | $a$ | $\tau (ns)$ | $\xi \left( \frac{amu}{ns} \right)$ |
| --- | --- | --- | --- | --- | --- |
| 2009-H1N1-ungly | 100% | $9.46 \times 10^{-6}$ | 1.15 | 47.1 | 438 |
| 2009-H1N1-gly | 83% | $5.90 \times 10^{-6}$ | 1.28 | 48.7 | 268 |
| 2009-H1N1-vir | 94% | $6.09 \times 10^{-6}$ | 1.09 | 51.5 | 313 |

Intuitively, glycosylation will increase the solvent accessibility of a protein by increasing its surface area, which we saw in our work (Figure SI3C). Increased solvent accessibility should result in a restriction of solvent motion; (58) we saw a corresponding, albeit small, restriction in solvent motion due to glycosylation (Table SI3). However, this was calculated including all the solvent around the protein, so most of the “signal” of the decrease in solvent motion due to glycosylation might have been lost in the noise. We carried out more detailed calculations honing in on how glycans affected the entropy and energy of the solvent through grid inhomogeneous solvation theory (see methods in SI (99-102)). We see that glycosylation reduces solvent entropy and orders the waters while reducing the amount of free water in the vicinity of the glycan (Table 3). Thus glycans can slow down water passage near the protein surface.

**Table 3.** Per-water thermodynamic properties of the solvent in the glycan-adjacent regions. Glycans restrict water movement and coordinate waters. The S_trans_ and S_orient_ are the translational and rotational entropies, respectively, in the protein frame of reference. More negative values indicate less entropy, thus showing that water is more constrained positionally and orientationally, respectively. Bulk water entropy values are set to be 0. The E_sw_ and E_ww_ are interaction energies per water molecule for solute-water and water-water interactions, respectively. More negative E_sw_ values indicate more favorable interactions between the water and the solute, and thus shows whether water is coordinating to the solute or not. These values would be more negative near charged side chains and less negative near nonpolar side chains. More negative E_ww_ values indicate more favorable interactions between water and the rest of the waters in the solvent. These values would be more negative where water is free and acting like bulk water, and less negative when water is sequestered. We averaged solvent entropy and energy around each of the glycans, and subtracted the entropy and energy from the same space on the 2009-H1N1-ungly construct from the 2009-H1N1-gly construct to arrive at a difference. The sample standard deviation is also presented.

| | $S_{trans} \left( \frac{kcal}{mol} \right)$ | $S_{orient} \left( \frac{kcal}{mol} \right)$ | $E_{sw} \left( \frac{kcal}{mol} \right)$ | $E_{ww} \left( \frac{kcal}{mol} \right)$ |
| --- | --- | --- | --- | --- |
| 2009-H1N1-ungly | $-0.05 \pm 0.04$ | $-0.06 \pm 0.04$ | $-0.48 \pm 0.31$ | $-8.89 \pm 2.26$ |
| 2009-H1N1-gly | $-0.35 \pm 0.10$ | $-0.36 \pm 0.09$ | $-1.89 \pm 0.54$ | $-2.11 \pm 0.48$ |
| difference | $-0.30 \pm 0.08$ | $-0.31 \pm 0.10$ | $-1.41 \pm 0.44$ | $6.78 \pm 1.83$ |

To decode the effects of glycans and lateral protein crowding on influenza NA, we examined three systems each differing by one variable: Two systems are identical except for the presence/absence of glycans, and two systems (including one of the two above) are identical except for the environment they were studied in. This allows us to do a direct comparison of the effect glycosylation and the membrane crowding has on protein stability and dynamics. We found that glycans generally increase the stability, stiffness and rigidity of NA while reducing fluctuations, entropy and the sampling of configuration space. Glycans also increase the degree of a large scale conformational change in NA. A crowded membrane protein environment increases the entropy of the glycan sequons and may promote a large scale conformational change. At the solvent level, glycans restrict water movement. Understanding how glycosylation and protein environment affect a protein’s characteristics informs the scenarios when it is acceptable to reduce the complexity of a protein model and still recover its native characteristics and when a reduction in complexity is not valid. There is ample space for future work to investigate how translatable these results are to other systems, and how well (or poorly) they hold up when creating larger protein environments that contain more realistic, biological environments.

## Supporting Information Available

Methods for system creation, running and analysis; Table SI1: the N-linked residues connecting each of the glycans to the protein; Figure SI1: protein rigidity, compactness and stability as a function of glycosylation and environment; Table SI2: rigidity of the systems as measured by RMSD clusters; Figure SI2: stability of severe acute respiratory syndrome (SARS) and SARS2 systems as a function of glycosylation; Figure SI3: protein fluctuations, entropy and solvent accessibility as a function of glycosylation; Figure SI4: dihedral entropy of the NA structure as a function of protein environment; Figure SI5: conformational flexibility and compactness of the NA system. Table SI3: effective viscosity of the solvent in the NA systems

## Acknowledgements

C.S. thanks Zied Gaieb for valuable mentorship, Christoph Bannwarth for valuable discussions on protein environment, J.C. Gumbart for sharing source data, Mike Gilson and Tom Kurtzmann for advice on using GIST, Stephen Wells and Dafydd Jones for access to FLEXOME and advice on using it, James Krieger for advice on ANM syntax, Pranav Khade for help with PACKMAN, Modesto Orozco and Adam Hospital for help with PCAsuite, and Berk Hess for advice on deriving stiffness parameters from principal components.

This material is based upon work supported by the National Science Foundation Graduate Research Fellowship Program under grant no. DGE-1650112 to C.S. This work was supported in part by the National Institutes of Health under grant no. T32EB009380 to C.S.

