## Supplementary Information for "Glycosylation and Crowded Membrane Effects on Influenza Neuraminidase Stability and Dynamics"

**Methods**

*NA setup and simulations*

The neuraminidase structure was extracted from previous work (1) and comes from the influenza A/swine/Shandong/N1/2009 (H1N1) strain. This structure was partially glycosylated, containing eight glycans on the tetramer. Monomer A contained one glycan, monomer B contained four glycans, monomer C contained one glycan, and monomer D contained two glycans. A sequon is the protein amino acid that is covalently bonded to the glycan; in influenza these are all asparagine residues. The N-linked residues for each of these glycans are listed in Table SI1. Due to the glycosylation asymmetry necessitated by the crowded virion environment, not every sequon was occupied. The exact structures of each glycan present are listed in Fig. S5 of the original model. (1)

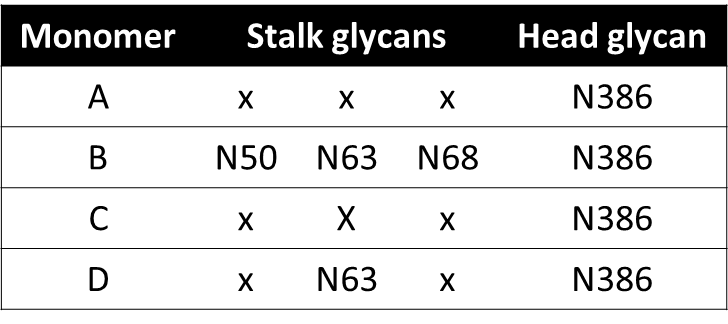

Table SI1. The N-linked residues connecting each of the glycans to the protein.

This structure was created and glycosylated in the virion context in previous work (1) so we will term it the 2009-H1N1-vir system. Before any of the MD simulations were run on the glycosylated virion, the NA structure and surrounding lipid bilayer was extracted for single protein simulations. From there, the glycosylated, single protein NA, which we will term the 2009-H1N1-gly system, was cloned and the glycans pruned to make an unglycosylated, single protein NA construct. We will call this last system the 2009-H1N1-ungly system.

These sets of simulations were then run: the single virion trajectory as described previously, (1) three replicates of the 2009-H1N1-gly system and three replicates of the 2009-H1N1-ungly system. The first set was the A/swine/Shandong/N1/2009 (H1N1) strain run in a whole virion environment previously. (1) In summary, this was run for 441.78 ns using NAMD 2.13 (2,3), with all the computational details described previously. (1) The second and third sets of simulations were run with essentially matching protocols to the whole virion simulation. For clarity, these single protein protocols are described here. They are described together as, after pruning the glycans, there were no differences in setting up or running the single protein simulations. The NA structure and lipid bilayer patch extracted from the virion environment was then sent through CHARMM-GUI (4-9) to generate resized periodic boundary conditions, new MD input files and solvent. The solvent consisted of explicit water molecules described through TIP3P (10) and 0.15 M NaCl described with an ion model (11) along with the catalytically important calcium ions. (12) CHARMM36 was used to parameterize the protein (13) along with the POPC lipid (14,15) and glycans. (16)

The 2009-H1N1-gly and 2009-H1N1-ungly systems were initially minimized with four alternating 250 step intervals of conjugate gradient (17) and adopted basis Newton-Raphson minimization (18) (and explained more fully here (19)), followed by a final 500-step interval. This totaled (4 x 250) + 500 = 1500 steps of conjugate gradient and 1500 steps of adopted basis Newton-Raphson minimization. Heating, in a traditional MD setup context, was not performed as it was not included in the default CHARMM-GUI workflow. Equilibration was then performed in three rounds of 25,000 steps each with a 1 fs timestep, progressively reducing the restraint force constants throughout the system. Next, the systems were equilibrated for another three rounds of 50,000 steps each with a 2 fs timestep while reducing the force constants further. The first two steps of equilibration used a three-step Langevin dynamics Verlet integrator (20) in the canonical (NVT) ensemble while the rest used a leapfrog Verlet integrator (21,22) in the isothermal-isobaric (NPT) ensemble with an Andersen-Hoover barostat (23,24) enforcing a pressure of 1 atm with a collision frequency of 20/ps. All equilibration steps used the Nosé–Hoover thermostat. (25-27) Periodic boundary conditions were created with a pair list generation of 16 Å, switching at 12 Å (28) and smoothing at 10 Å.

For the production MD, the barostat for all steps was a semi-isotropic combination of the Nosé–Hoover pressure control method (23) with the piston fluctuation control implemented with Langevin dynamics; (29) this is the standard barostat in NAMD simulations. Constant temperature was enforced through a Langevin temperature coupling thermostat. (30) Simulations were run in NPT at a pressure of 1 atm and a collision frequency of 50/ps with the Brünger−Brooks−Karplus integrator, (31) which is an extension of the Verlet algorithm. Atomic pair lists were created at 16 Å with a non-bonded cutoff at 12 Å and switching at 10 Å. (28) A timestep of 2 fs was used in the production MD simulations, which were run for 450 ns but results are presented for the first 441 ns to match closely with the virion simulation. Frames were saved every 10 ps.

The SHAKE algorithm (32) was used to constrain bonds involving hydrogen and long range electrostatic interactions were treated with particle mesh Ewald (33) in both the CHARMM equilibration and the NAMD production MD. The simulations were run at the Texas advanced computing center’s flagship supercomputer, Stampede2 – a member of the Extreme Science and Engineering Discovery Environment (34) ecosystem. PDB and PSF files from the simulations that were used for analysis are hosted on Github (https://github.com/cgseitz), while the larger trajectory files can be provided upon request.

*Protein rigidity*

We measured protein rigidity through the FLEXOME software program, (35) adopting methods seen previously. (36) FLEXOME is available on request from the University of Bath (<https://doi.org/10.15125/BATH-00940>). Only the α-carbons were analyzed as these should be the most rigid parts of the protein, and thus changes in their rigidity should be indicative of global changes.

Rigidity was also measured using the Lindemann coefficient (37) through the PCAsuite program (accessible here: https://mmb.irbbarcelona.org/software/pcasuite/), which is integrated into the pyPCcazip program. (38) Trajectories were zipped with 90% quality using only the α-carbons as the software did not contain enough memory to handle the full structure. Input and output files for the FLEXOME analysis are hosted on Github (https://github.com/cgseitz).

*Clustering*

The 2009-H1N1-gly trajectories were then concatenated; the same was done for the 2009-H1N1-ungly trajectories. These long trajectories, along with the single replicate of the 2009-H1N1-vir system, were then clustered using GROMACS-based GROMOS clustering. (39) Clustering was performed with an RMSD cutoff of 0.5 Å, 0.6 Å, 0.7 Å, 0.8 Å, 0.9 Å and 1.0 Å and the number of resulting clusters was calculated as a percentage of the total number of frames. This adapted a method seen previously. (40) Trajectory files used for clustering can be provided on request, while the rest of the input and output clustering files are hosted on Github (https://github.com/cgseitz).

*RMSD*

Root mean square deviations (RMSD) of the MD trajectories were calculated through MDTraj. (41) The trajectories of all systems were aligned by their α-carbons before their RMSD were calculated. This was then calculated separately for the NA head and NA stalk to focus on differences special to those regions. The NA head RMSD was calculated after aligning all simulations to the NA head alpha carbons in the first frame of their respective simulations, while the NA stalk RMSD was calculated after aligning all simulations to the NA stalk alpha carbons in the first frame of their simulations. The standard deviation was also calculated.

Separately, RMSD was also calculated through MDTraj for the severe acute respiratory syndrome (SARS) SARS-CoV and SARS-CoV-2 spike glycoprotein receptor binding motif in complex with the angiotensin-converting enzyme 2 protein. These simulations were run previously (42) with the source data generously provided by the authors. A simple Jupyter notebook detailing the RMSD work is hosted on Github (https://github.com/cgseitz) while the aligned input trajectories can be provided upon request.

*Principal component analysis*

We used principal component analysis (PCA) to analyze the conformational space accessed by the simulations. We built off a method described previously. (43) This involved creating each of the three systems’ simulations in the same PCA space and plotting each system’s first two principal components in that space as a function of free energy. For example, we combined all three replicates of the 2009-H1N1-gly system together and plotted them together in the same PCA space as the other systems to allow for direct comparisons. We then measured the 2D area that each system covered and divided by the number of replicates of each system, i.e. three for the 2009-H1N1-gly and 2009-H1N1-ungly systems and one for the 2009-H1N1-vir system. The 2009-H1N1-ungly systems traversed the most PC space, so we set that space to be “100% area covered” and calculated how much relative space the 2009-H1N1-gly simulations covered and how much relative space the 2009-H1N1-vir trajectory covered. Although these simulations were all set in the same PC space and they covered similar areas, it is useful to note that there is not a full overlap between them. For example, though we report that the 2009-H1N1-vir trajectory covered 94% of the space as the three 2009-H1N1-ungly replicates, that does not mean the three 2009-H1N1-ungly replicates covered all of the space that the 2009-H1N1-vir replicate covered: this is simply a method of comparing total areas sampled with each other.

To measure the stiffness/depth of the PCA space accessed by each system we examined the eigenvalues and their corresponding force constants of the free energy spaces. This was modeled after previous methods. (44-46) We calculated the first principal component for each system along with its corresponding eigenvalue and used that to derive the force constant defining the harmonic well in the direction of the first principal component. The eigenvalue is a measure of the stiffness of the system while the force constant is a measure of the depth and sharpness of the free energy wells. This can be calculated as follows: $k= \frac{k_{B}T}{100\cdot\lambda}$ where $k$ is the force constant we are looking for, $k_{B}$ is the Boltzmann, equivalent to 8.314462618 x 10^-3^ $\frac{kJ}{mol\cdot K}$ and $\lambda$ is the eigenvalue derived from the principal component. The MD trajectories were run with coordinate data in Å which means the corresponding eigenvalues of the principal components would be in units of Å^2^. Thus we divided the eigenvalues by 100 so they would be in units of nm^2^ for a more standard force constant value. After calculating the force constant for each replicate of the 2009-H1N1-gly and 2009-H1N1-ungly simulations we averaged them together. The depth of the free energy space was calculated as follows. We first calculated the autocorrelation function of the first principal components, which were previously derived as described above. We fitted that data and the time length of the simulations (~440 ns) to a two-variable exponential function $y=ae^{\frac{-t}{\tau}}$ where $y$ is the autocorrelation function and $t$ is the time length of the simulations. This fitting gave us $a$ and $\tau$, where $\tau$ is then used to calculate the free energy well depth $\xi$: $\xi=k\cdot\tau$. Note that $\xi$ does not exactly equal the $k\cdot\tau$ presented in the results here with the given significant figures as full decimal places were carried through in the calculations before being rounded at the end, as shown below. The units conversion is not trivial so we will give an example calculation here, using the knowledge that $amu=\frac{g}{mol}$ and that $J=\frac{kg\cdot m^{2}}{s^{2}}$.

$9.46 x {10}^{-6}\frac{kJ}{mol\cdot{nm}^{2}}\cdot47.1 ns\cdot\frac{mol\cdot amu}{g}\cdot\frac{kg\cdot m^{2}}{J\cdot s^{2}}\cdot\frac{1000 J}{1 kJ}\cdot\frac{1 x {10}^{9} nm}{1 m}\cdot\frac{1 x {10}^{9} nm}{1 m}\cdot\frac{1000 g}{1 kg}\cdot\frac{1 s}{1 x {10}^{9} ns}\cdot\frac{1 s}{1 x {10}^{9} ns}=445\frac{amu}{ns}$. This is close, but not equivalent, to our presented result of 438 $\frac{amu}{ns}$ as our presented results carried forward full decimal places for each quantity until being rounded at the end. A Jupyter notebook going through the PCA steps is hosted on Github (https://github.com/cgseitz) while the trajectory files can be provided.

*Elastic network models*

Gaussian network models (GNM) (47-49) and anisotropic network models (ANM) (50) were created with the ProDy software package (51,52) using code adapted from previous work. (53) For simplicity we looked at the node with the lowest frequency only, as this gives some sense of the system dynamics (54) for both the GNM and ANM work. The Kirchoff matrix in the GNM was built with a pairwise interaction cutoff distance of 10 Å and the Hessian matrix in the ANM was built with a pairwise interaction cutoff distance of 15 Å and a spring constant of 1.0. The eigenvalues were calculated using only the α-carbons as the input, as these should be the most stable parts of the protein. Thus if these are moving, they should show global trends, not small side chain fluctuations that are not important to large scale conformational changes. The elastic network model analysis is hosted on Github (https://github.com/cgseitz).

*RMSF*

Root mean square fluctuations (RMSF) of the 2009-H1N1-gly and the 2009-H1N1-ungly systems were calculated through CPPTraj. (55) The structures were aligned by their α-carbons. With the four NA head N-linked sequons determined (details in the system preparation methods section), RMSF calculated were made through shells of increasing distance through CPPTraj on both the 2009-H1N1-gly and 2009-H1N1-ungly systems. RMSF was calculated for all residues within 5 Å of each NA head sequon, 10 Å, 15 Å, 20 Å, 25 Å, and lastly 30 Å. Each shell was then averaged together across all replicates of the system for 12 RMSF calculations at each distance: 1 distance selection x 4 homomonomers/NA x 3 MD replicates = 12 RMSF calculations. At each distance, the RMSF values coming from the 2009-H1N1-gly systems were subtracted from those coming from 2009-H1N1-ungly systems to see how much higher in RMSF, and thus how much more flexible, the unglycosylated constructs were at each distance. The standard deviation was also calculated. Simple scripts for the RMSF can be found in a Jupyter notebook on Github (https://github.com/cgseitz).

*Entropy*

The water entropy was measured through Amber’s implementation of the grid-based inhomogeneous solvation theory (GIST), (56,57) specifically the GPU-accelerated version. (58) A glycosylated NA structure and a corresponding unglycosylated structure were selected from the MD simulation inputs, described previously. These systems were put through six rounds of equilibration steps, detailed here. Rounds (1) and (2) were rounds of minimization, rounds (3) and (4) were rounds of heating, and rounds (5) and (6) were rounds of full equilibration. Minimization used 400 ps in each round. Round (1) minimized the waters while restraining the protein while round (2) minimized the waters and protein hydrogen atoms. Round (3) heated the system from 0 K to 50 K while restraining the protein heavy atoms. Round (4) heated the system from 50 K to 300 K; each round of heating lasted 200 ps. Round (5) lasted 10 ns while round (6) lasted 5 ns. Rounds (3) – (6) used SHAKE(32) and a Langevin thermostat(30) with a collision frequency of 2/ps. Rounds (3) and (6) were in the NVT ensemble, while rounds (4) and (5) were in the NPT ensemble using the Monte Carlo barostat. (59) All rounds used the steepest descent method followed by a conjugate gradient (17) and the leapfrog Verlet integrator of motion. (21,22) After the equilibration, the systems were run for 100 ns of full MD using Amber 18 (60,61) with the NPT ensemble. The GIST properties were then calculated with through gistpp (62) for each glycan. One calculation grid was centered around each glycan using the FindCentroid.py script provided with GIST. These coordinates were then used for the unglycosylated construct to make a direct comparison of how the glycans are affecting water properties. The glycan volume was then subtracted from the calculation grid while retaining a 3 Å water shell around the heavy atoms of the glycans per the GIST documentation, for both the glycosylated and unglycosylated constructs. The water translational entropy (dTS_trans_), water orientational entropy (dTS_orient_), mean water-solute interaction energy (E_sw_), and mean water-water interaction energy (E_ww_) were then measured for each glycan region and then averaged amongst all eight glycan regions. We then subtracted the unglycosylated construct values from the glycosylated construct values to find the difference. Then the per-water dTS_trans_, dTS_orient_, E­_sw_, and E_ww_ were determined by calculating the volume of the water shell in the calculations above, and multiplying that times the average entropy or energy per voxel and multiplying that times bulk water density. This was done as there may be a slightly different number of waters between the glycosylated and unglycosylated systems, so summing the total energies may produce misleading results.

Protein conformational “packing” entropy was measured through the packing entropy webserver. (63) Structures from the MD simulation inputs were used: One glycosylated structure and one unglycosylated structure. These files were then modified to contain the element for each atom, per PDB format, so the server would accept the files. The surface probe radius was 1.4 Å, and the number of points generated around each point used for the surface was 30. After the calculations completed, the entropy of the N-linked sequons was extracted for comparison.

Conformational entropy of the φ- and φ-angles were measured through the X-Entropy software program.(64) To use this, we calculated the dihedral angles, specifically the φ- and φ-angles using MDTraj (41) for each system’s MD trajectories: 2009-H1N1-gly, 2009-H1N1-ungly, and 2009-H1N1-vir. We then averaged the angle values for the 2009-H1N1-gly replicates together, and averaged the angle values for the 2009-H1N1-ungly replicates together. Finally we wrote a python script to measure the angle entropy in shells, up to n=5, around each N-linked glycan. For example, N891 is one of the N-linked sequons. The n=1 shell averaged the entropy values from residues 890, 891 and 892 (± 1 from the N-linked sequon). Similarly, the n=2 shell contains the residues that are ± 2 residues from each sequon, and so on up to n=5. We did this to examine a distance dependence for the entropy results seen. The sample standard deviation was also calculated from the entropy difference at each of the eight sequons. Most of the entropy input and output files can be found on Github (https://github.com/cgseitz); however, the trajectories are not hosted there and can be provided on request.

*Head tilt*

The method for the NA head tilt calculation was published previously. (1) In summary, this involved selecting three different points in the NA structure and calculating how this angle changed over the course of the simulations. The first point is the center of mass of the α-carbons of the upper stalk region (residues 50 to 78), the second point is the α-carbon of residue 78 (a hinge residue between the stalk and the head), and the third point is the center of mass of the α-carbons of the head region (residues 91 to 469). We also performed this calculation on some previously simulated NA tetramers (65) for comparison. The standard deviation was also calculated. A TCL script calculating the head tilt can be found on Github (https://github.com/cgseitz).

*Radius of gyration*

The radius of gyration was calculated with MDTraj. (41) To plot all the simulations on the same space, the 2009-H1N1-gly and 2009-H1N1-ungly simulations were set with a stride of 25 and the 2009-H1N1-vir trajectory was set with a stride of 4. The 2009-H1N1-gly trajectories were then averaged together before plotting, and the same was done with the 2009-H1N1-ungly trajectories. We calculated the radius of gyration both with and without the glycans for the 2009-H1N1-gly and 2009-H1N1-vir systems, and present them separately. The standard deviation was also calculated. Tidy scripts detailing the radius of gyration calculations are found on Github (https://github.com/cgseitz).

*Local viscosity*

Local viscosity was calculated for the oxygens present in the simulations’ water solvent, and also for the NA in the simulations. This was done for all systems; the 2009-H1N1-gly systems were then averaged together and the 2009-H1N1-ungly systems were treated the same way. The method of calculating local water and protein viscosity is the same so they will be described together here. To start, the diffusion constant (whether it be for water or for protein) was calculated for the simulation of interest through CPPTraj. (55) The viscosity can then be calculated from the diffusion constant: $D=\frac{k_{B}T}{6\pi\eta r}$ where $D$ is the diffusion constant we just calculated, $k_{B}$ is the Boltzmann constant set at 1.380649 x 10^-23^ $\frac{m^{2}kg}{s^{2}K}$, $T$ is the simulation temperature 298 K, $\eta$ is the viscosity we are looking for, and $r$ is the radius of the particle we are examining. The radius for oxygen is 1.7682 Å (taken from the TIP3P water model). The protein is not a sphere so an exact radius cannot be determined; as we are interested in diffusion through the membrane we measured the transmembrane stalk region of NA to have a radius of ~10 Å. We could not calculate the effective water viscosity in the 2009-H1N1-vir system because the individual solvent particles keep moving in and out of the protein’s frame of reference and would not necessarily stay localized to the protein. This is not an issue in the 2009-H1N1-gly or 2009-H1N1-ungly simulations as periodic boundary conditions mean each solvent particle must keep interacting with the protein, which allows a diffusion constant to be calculated. The trajectories used for this analysis can be sent upon request, while the simple scripts to calculate the local viscosity are hosted on Github (https://github.com/cgseitz).

*Solvent accessible surface area*

The solvent accessible surface area (SASA) was calculated through CPPTraj (55) using the linear combinations of pairwise overlaps method. (66) The 2009-H1N1-gly and 2009-H1N1-ungly SASA was computed with a stride of six to make for more direct comparisons with the 2009-H1N1-vir system, as that dataset was created with six times fewer frames being saved out to disk. The scripts used for calculating the SASA are hosted on Github (https://github.com/cgseitz).

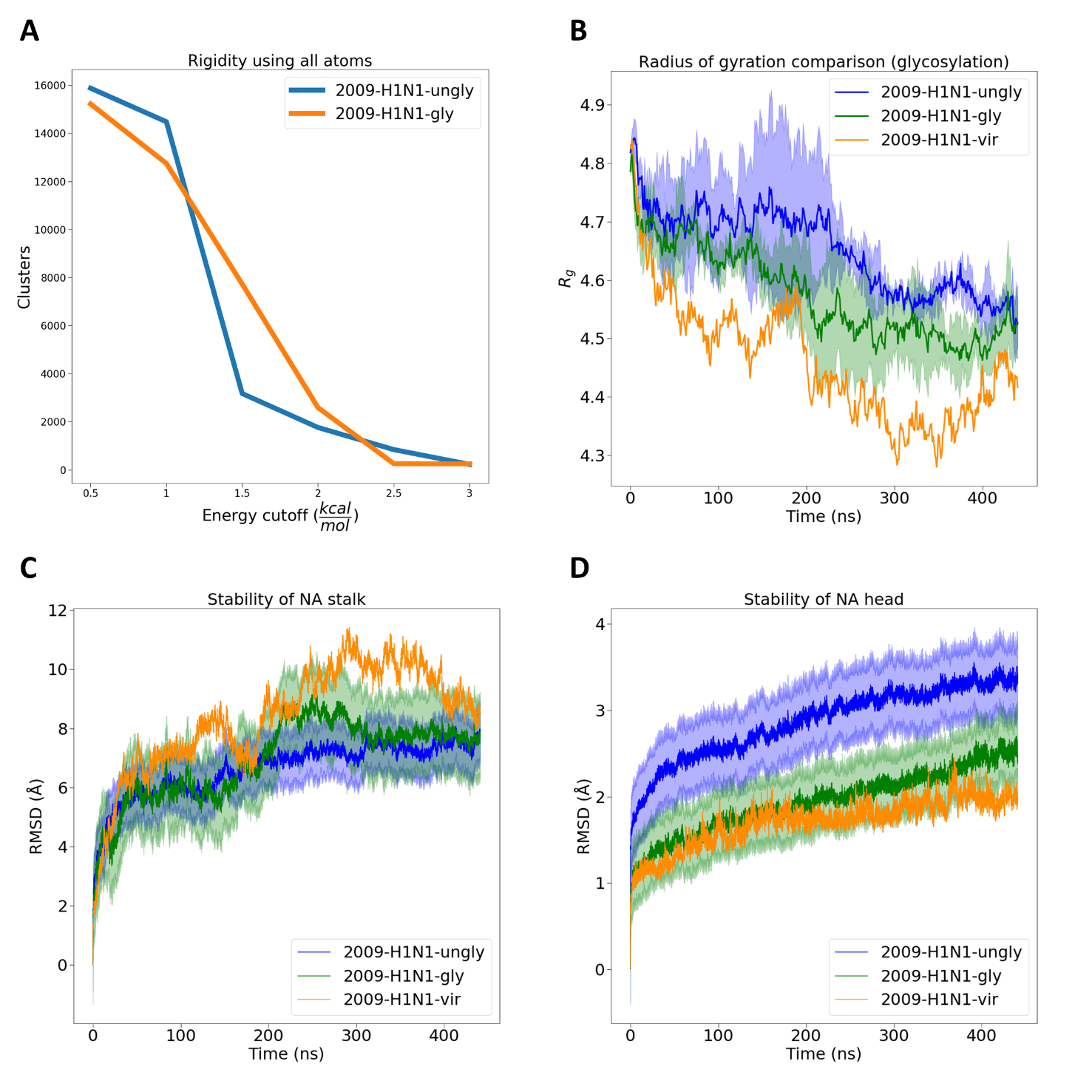

Figure SI1. Protein rigidity, compactness and stability as a function of glycosylation and environment. (A) Rigidity of the NA systems computed using all atoms. This shows the number of clusters that retain nonbonded interactions at different energy cutoffs, and shows the same trends as seen in Figure 2A: the 2009-H1N1-ungly system initially is more rigid before the 2009-H1N1-gly system becomes more rigid at slightly higher energy cutoffs. (B) Radius of gyration of the NA systems, with no glycans included in the calculations. The 2009-H1N1-ungly and 2009-H1N1-gly systems have the standard deviation of the *R_g_* calculations shaded. (C) NA stalk stability. The 2009-H1N1-ungly and 2009-H1N1-gly systems have the standard deviation of the RMSD calculations shaded. (D) NA head stability. The 2009-H1N1-ungly and 2009-H1N1-gly systems have the standard deviation of the RMSD calculations shaded.

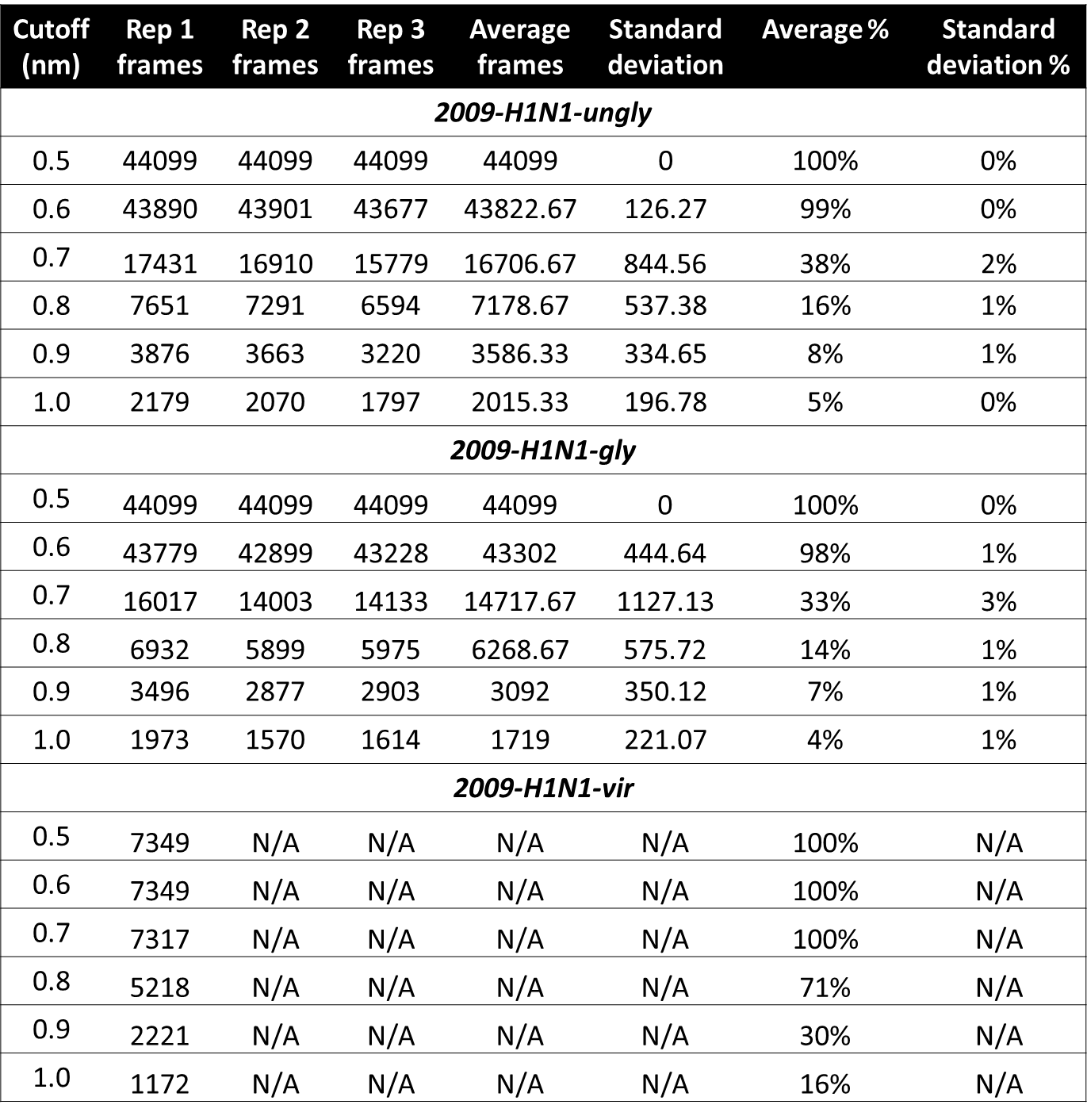

Table SI2. Rigidity of the systems as measured by RMSD clusters. The number of clusters found at each distance cutoff was then divided by the total number of frames (and thus the maximum number of clusters) to see how many clusters were formed out of a total possible 100% of frames being unique clusters.

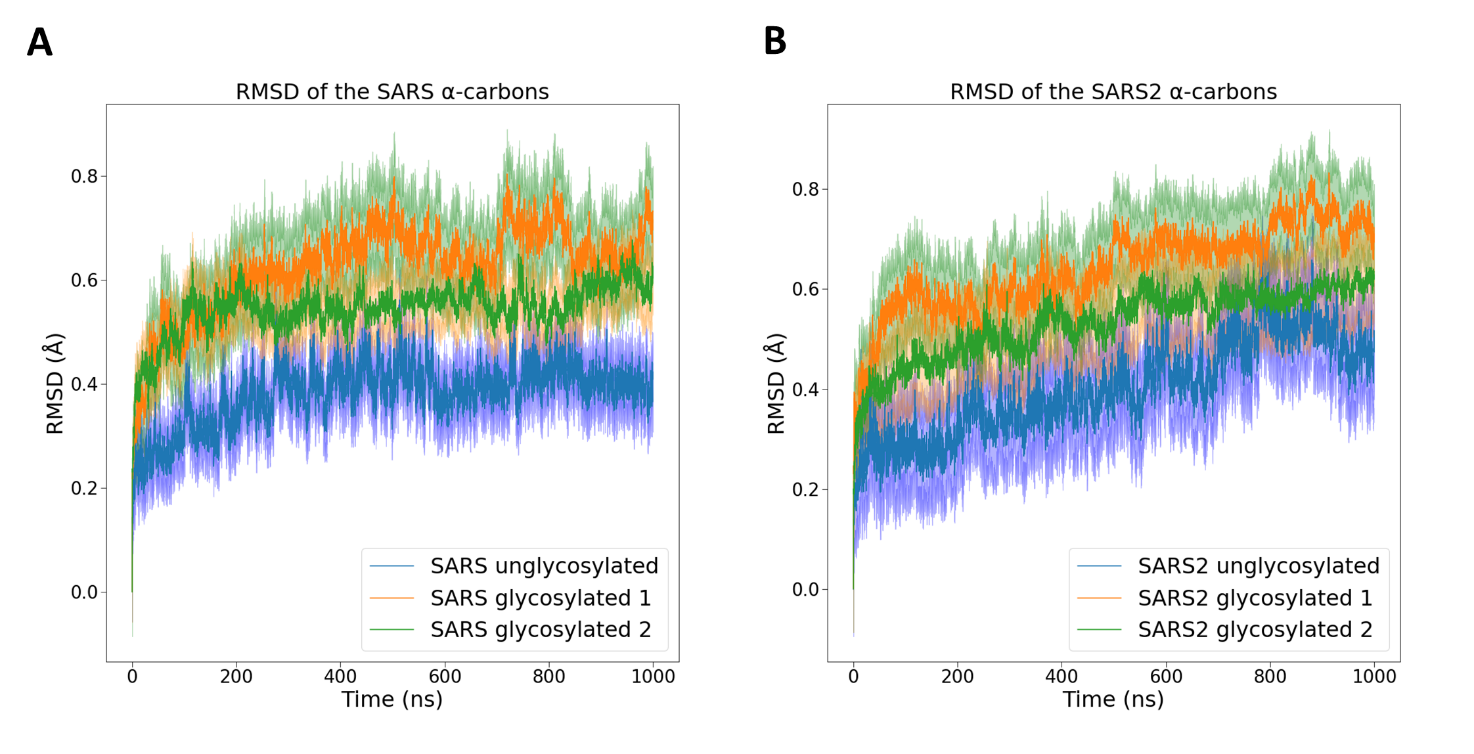

Figure SI2. Stability of severe acute respiratory syndrome (SARS) and SARS2 systems as a function of glycosylation. Each plot, (A) and (B), includes one unglycosylated system and two glycosylated systems with different glycoprofiles, with the standard deviations shaded. Glycans increase the stability across systems. These systems come from previously published work. (42)

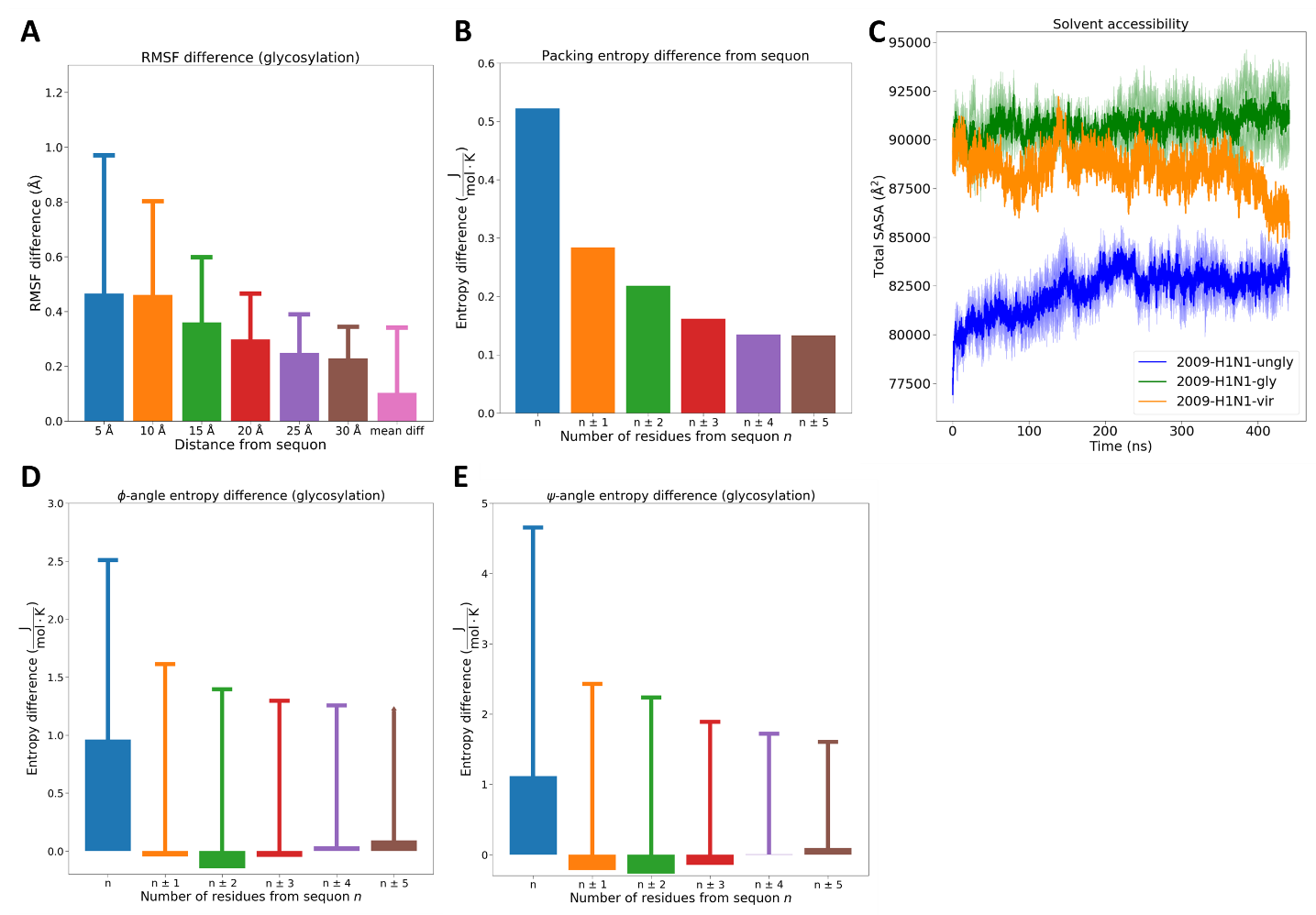

Figure SI3. Protein fluctuations, entropy and solvent accessibility as a function of glycosylation. (A) RMSF calculations of the NA head. Subtracting the RMSF values of the 2009-H1N1-gly system from the 2009-H1N1-ungly system results in the “RMSF difference” shown in the figure. In other words, this figure shows how much larger the RMSF values of the 2009-H1N1-ungly system are than the RMSF values of the 2009-H1N1-gly system. (B) Packing entropy of the NA structure. Subtracting the 2009-H1N1-gly entropy from the 2009-H1N1-ungly entropy we arrive at an “entropy difference” which was measured for up to five residues before and after the sequon, with each “sequon” here being an average from all eight occupied sequons in each system. For reference, the average packing entropy difference across the entire protein is 0.0525 $\frac{J}{mol\cdot K}$. (C) SASA of the NA systems. Glycans increase the SASA of the system while a crowded environment decreases it. (D) and (E) Dihedral entropy of the NA structure as a function of glycosylation. We measured the φ (D) and ψ (E) angles in the 2009-H1N1-gly and 2009-H1N1-ungly systems. We then subtracted the 2009-H1N1-gly entropy from the 2009-H1N1-ungly entropy to arrive at an “entropy difference”, again using an average of all eight occupied sequons in each system. The average difference for the φ-angle is 0.175 $\frac{J}{mol\cdot K}$, and the average difference for the ψ-angle is 0.170 $\frac{J}{mol\cdot K}$.

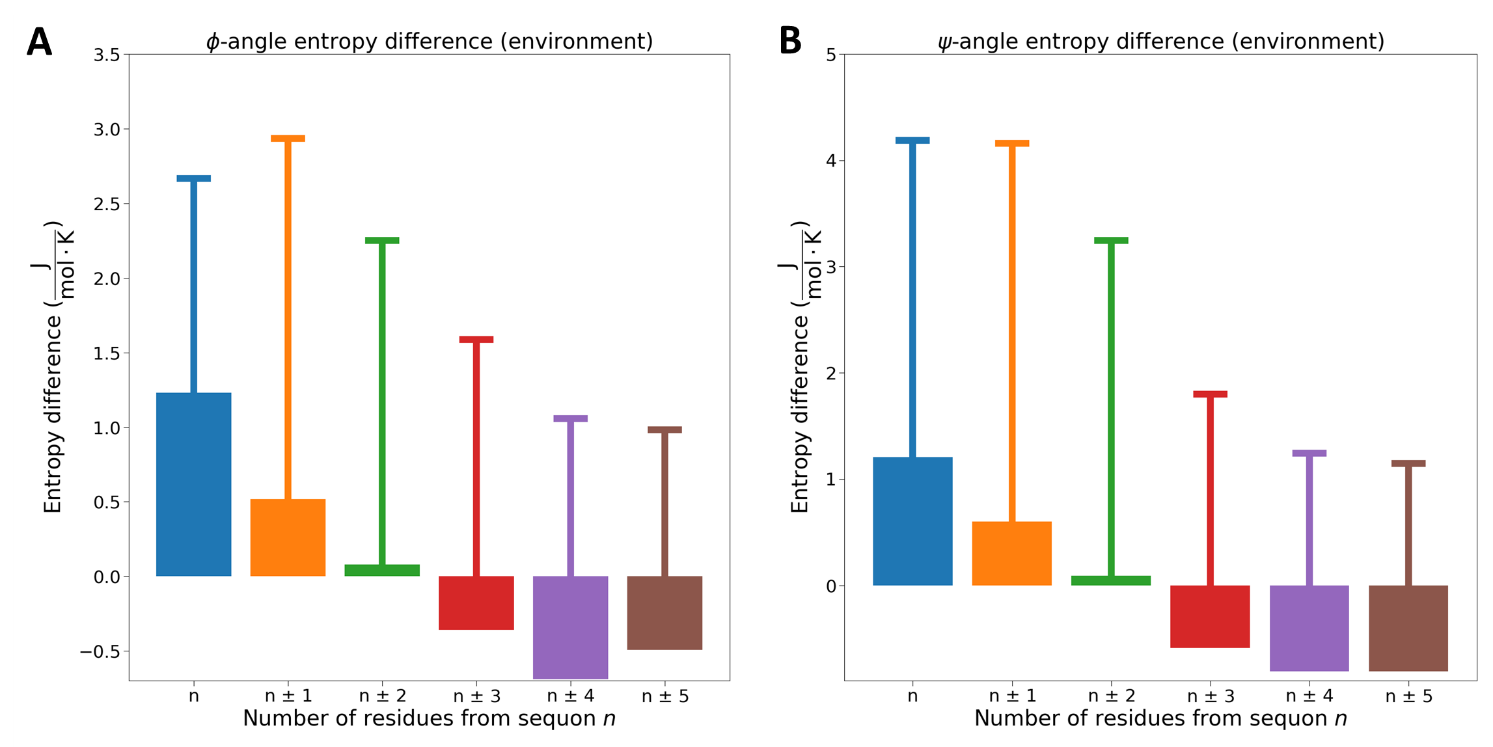

Figure SI4. Dihedral entropy of the NA structure as a function of protein environment. We measured the φ and ψ angles in the 2009-H1N1-gly and 2009-H1N1-vir systems. We then subtracted the 2009-H1N1-gly entropy from the 2009-H1N1-vir entropy to arrive at an “entropy difference”. We averaged entropy results from all eight occupied sequons in each system to arrive at the results here. The average difference for the φ-angle is -0.116 $\frac{J}{mol\cdot K}$, and the average difference for the ψ-angle is -0.137 $\frac{J}{mol\cdot K}$.

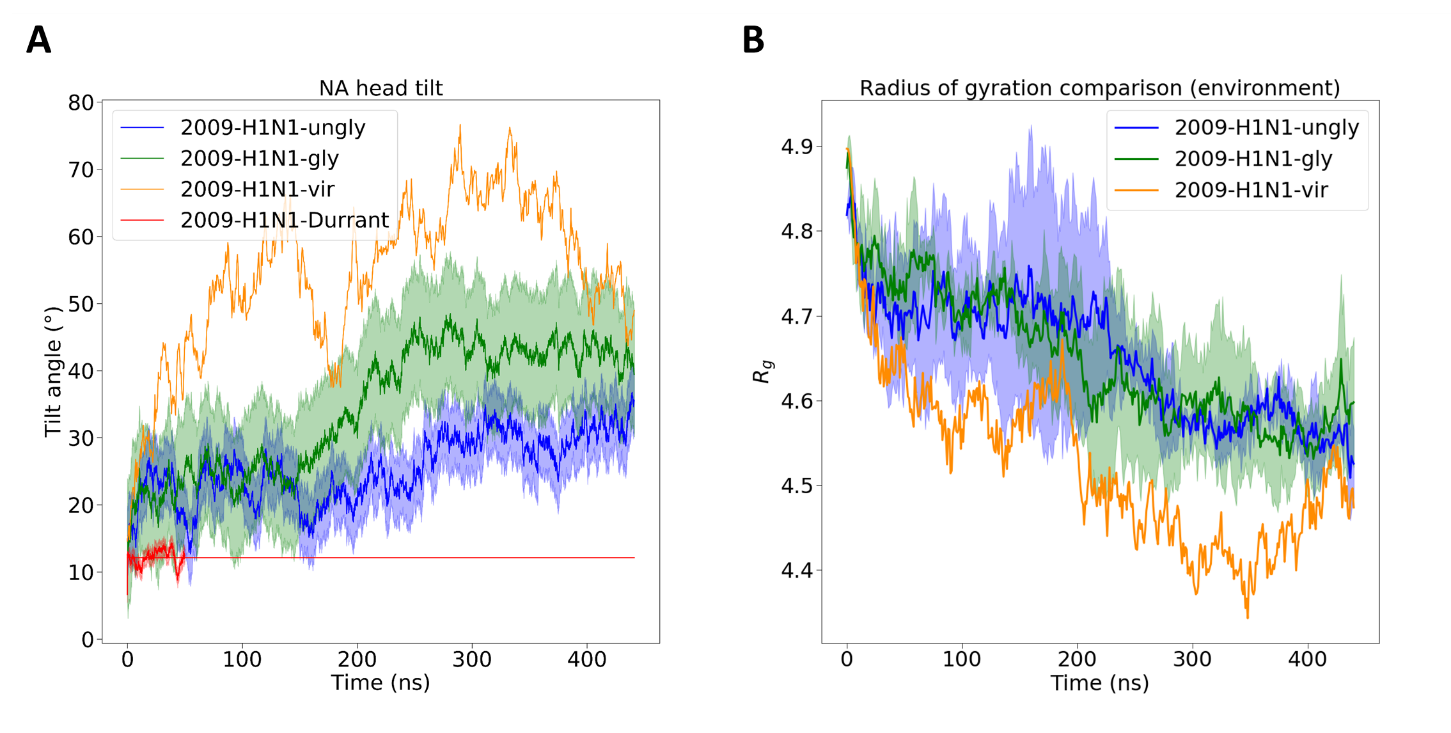

Figure SI5. Conformational flexibility and compactness of the NA system. (A) NA head tilting. The 2009-H1N1-Durrant system consists of 50 ns simulations with an unglycosylated NA and was published previously; (65) we have plotted the NA head tilting in that system and extended a straight line representing the mean tilt. The 2009-H1N1-ungly, 2009-H1N1-gly, and 2009-H1N1-Durrant systems have the standard deviation of the tilt angle calculations shaded. (B) Radius of gyration of the NA systems, with glycans included in the calculations of the 2009-H1N1-gly and 2009-H1N1-vir systems. The 2009-H1N1-ungly and 2009-H1N1-gly systems have the standard deviation of the *R_g_* calculations shaded.

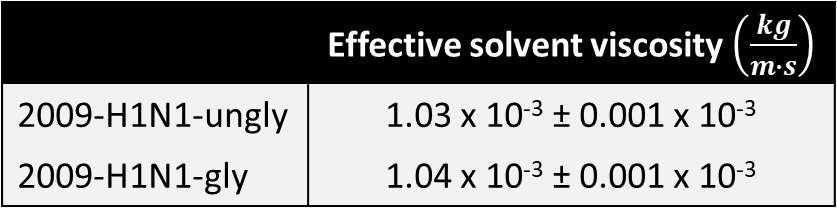

Table SI3. Effective viscosity of the solvent in the NA systems. Glycans increase the effective viscosity of the solvent. The effect is relatively small as the entire solvent was considered. The sample standard deviation is also presented.
